# Feasibility of Using Wearable EMG Armbands combined with Unsupervised Transfer Learning for Seamless Myoelectric Control

**DOI:** 10.1101/2022.01.06.475232

**Authors:** M. Hongchul Sohn, Sonia Yuxiao Lai, Matthew L. Elwin, Julius P.A. Dewald

## Abstract

Myoelectric control uses electromyography (EMG) signals as human-originated input to enable intuitive interfaces with machines. As such, recent rehabilitation robotics employs myoelectric control to autonomously classify user intent or operation mode using machine learning. However, performance in such applications inherently suffers from the non-stationarity of EMG signals across measurement conditions. Current laboratory-based solutions rely on careful, time-consuming control of the recordings or periodic recalibration, impeding real-world deployment. We propose that robust yet seamless myoelectric control can be achieved using a low-end, easy-to-“don” and “doff” wearable EMG sensor combined with unsupervised transfer learning. Here, we test the feasibility of one such application using a consumer-grade sensor (Myo armband, 8 EMG channels @ 200 Hz) for gesture classification across measurement conditions using an existing dataset: 5 users x 10 days x 3 sensor locations. Specifically, we first train a deep neural network using Temporal-Spatial Descriptors (TSD) with labeled source data from any particular user, day, or location. We then apply the Self-Calibrating Asynchronous Domain Adversarial Neural Network (SCADANN), which automatically adjusts the trained TSD to improve classification performance for unlabeled target data from a different user, day, or sensor location. Compared to the original TSD, SCADANN improves accuracy by 12±5.2% (avg±sd), 9.6±5.0%, and 8.6±3.3% across all possible user-to-user, day-to-day, and location-to-location cases, respectively. In one best-case scenario, accuracy improves by 26% (from 67% to 93%), whereas sometimes the gain is modest (e.g., from 76% to 78%). We also show that the performance of transfer learning can be improved by using a “better” model trained with “good” (e.g., incremental) source data. We postulate that the proposed approach is feasible and promising and can be further tailored for seamless myoelectric control of powered prosthetics or exoskeletons.

## 1. INTRODUCTION

Myoelectric control uses electromyography (EMG) signals as human-originated input to enable intuitive interfaces with machines. EMG signals, often recorded transcutaneously, pick up an ensemble of motor neuron action potentials representing neural commands from the brain to control movement. Thus, by decoding these signals the movement goal of a person can be inferred [1]. As such, many human-machine interface applications employ myoelectric control to autonomously predict user intent or operation mode using machine learning based classifiers such as k-Nearest Neighbors, Linear Discriminant Analysis, or Support Vector Machines [2–4], and more recently using deep learning models such as Convolutional Neural Networks (CNN) or Recurrent Neural Networks [5, 6]. Such online prediction models can be used, for example, in assistive technologies such as powered prosthetics [4, 7, 8], when interacting with robotic devices [9–11], or controlling a virtual avatar [12–14].

However, performance of myoelectric control-based prediction models inherently suffers from the non-stationarity of EMG signals: the magnitude, frequency, and spatial characteristics of the signals vary across measurement conditions [15, 16]. Sources of such variation include noise, skin conductivity, sensor placement, and physiological differences between people, leading to performance differences between different users or even the same user at different times [4]. Variations in input EMG signals used for training vs. testing, or even within training result in degraded performance (e.g., prediction accuracy) [17]. One of the most common laboratory-based solutions is to retrain or periodically calibrate the model with new data, either for each user or for each session [18]. However, such procedures require additional data collection and testing/validation of the model, which is time-consuming and costly, and thus impedes real-world deployment.

Another practical limitation in translating myoelectric control system from the lab to the real-world is that current development largely relies on high-end, research-grade sensors. Careful recording (e.g., proper preparation and sensor placement) of high-quality EMG data (e.g., high signal-to-noise ratio, sampling rate or bandwidth, and resolution) and increased numbers of channels (e.g., multi-electrode arrays) improve robustness and performance [19, 20]. However, these benefits come at the cost of increased expense and time for data collection, greater power consumption and computational overhead. Furthermore, such advanced systems are not readily adopted in the field especially in medical applications, where simplicity and intuitive usability (e.g., by patients and/or clinicians) are crucial factors for technology acceptance [21–23].

We propose that robust yet seamless myoelectric control can be achieved using low-end, easy-to-“don” and “doff” wearable sensors combined with unsupervised transfer learning. Consumer-grade wearable sensors offer many advantages over laboratory-grade equipment including affordable cost, extended recording time (low power), and streamlined user applicability [21, 22]. On the other hand, transfer learning leverages techniques to generalize a model pretrained with an initial source data to newly recorded target data with minimal or no recalibration [24]. For example, various domain adaptation or generalization algorithms [25] have been proposed to improve prediction accuracy across different users, sessions, or sensor locations [26–28]. Such combinations remove the burden from users, enabling them to just wear the sensor. To our knowledge, however, no study has systematically investigated whether, and in what contexts unsupervised learning can be applied to various EMG measurement conditions (such as users, sensor locations, and times) without losing performance, especially when the data is collected from low-end wearable sensors. [12, 20].

Here, we test the feasibility of one such application using a consumer-grade armband sensor for robust gesture classification across various measurement conditions. Using an existing dataset [29] and by adopting an unsupervised transfer learning algorithm developed by Côté-Allard et al. [30], we demonstrate that the proposed approach, which automatically adjusts any deep learning model trained with source data (i.e., data from a particular user, day, or sensor location) to improve classification performance for unlabeled target data for a different user, day, or location, is feasible and promising for enabling seamless myoelectric control.

## 2. MATERIALS AND METHODS

### 2.1. EMG dataset and wearable sensor

We used publicly available dataset [29] with surface EMG signals recorded from 5 users × 10 days × 3 sensor locations during various wrist and hand gestures. Detailed description can be found in the original paper [29] and corresponding data repository. Briefly, five young healthy individuals participated. Data was acquired using the wearable sensor Myo Armband (Thalmic Labs, Kitchener, Canada; Fig. 1). The device can record 8 channels of EMG from equally spaced dry electrodes, which allows users to simply slip the sensor onto the forearm without any skin preparation. Importantly, such bracelet type sensor has been reported to be the most acceptable from among many wearable sensors [21–23].

**Figure 1.**
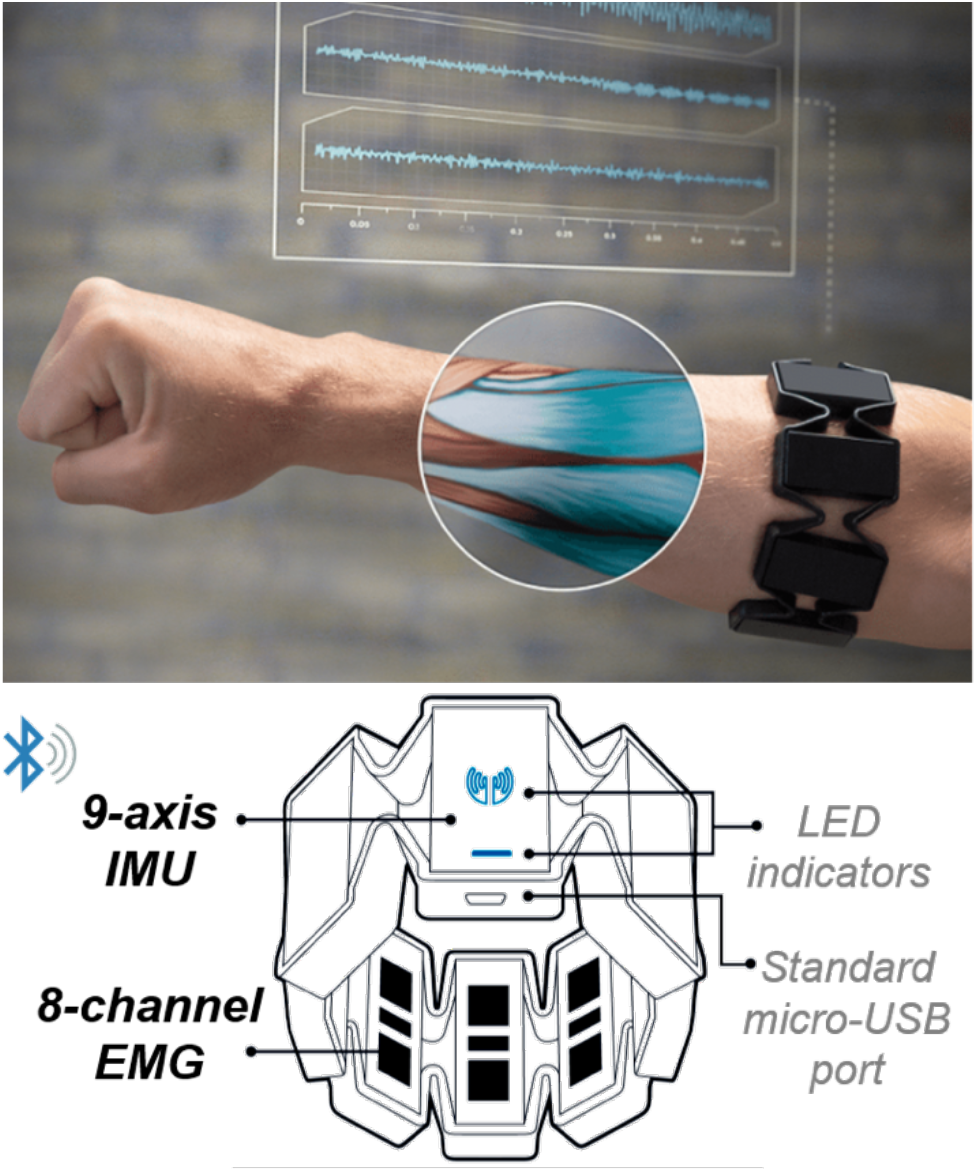
Consumer-grade wearable EMG sensor Myo Armband (Thalmic Labs, Kitchener, Canada). Typical sensor location on forearm (top) and detailed components (bottom): 9-axis IMU and 8-channel EMG sampled at 50 Hz and 200 Hz, respectively, communicated via BLE.

Measurements took place for multiple days where on each day, users wore Myo Armband at one of three different locations: Neutral position (N), 8-mm anticlockwise inward rotation (I), and 8-mm clockwise outward rotation (O). Measurements from each location were acquired 10 times over 30 consecutive days on random permutation order with uniform distribution.

Each trial consisted of 1.5-s data from 8 channels of EMG that encapsulated the onset and steady phase of muscle contraction during one of 22 gestures, comprised of various combinations of wrist and hand (eight 1-DoF and fourteen 2-DoF) motions [3]. Each gesture was repeated 4 times in each day. EMG signals were sampled at 200 Hz and high-pass filtered at 15 Hz with a fifth-order Butterworth filter. We further segmented each 1.5-s length data using sliding window of 250 ms (with 80% overlap) for feature extraction (see section 2.2.2 below).

In summary, EMG data structure from each user, at any day, at any sensor location, was shaped as 4 × 22 × 26 × 8 (repetitions × gestures × segments × channels) and was used as input for deep learning models (see below). This dataset will be now referred to as the Myo Dataset.

### 2.2. Transfer learning framework

To test the feasibility of preserving robust performance of myoelectric control system in the context of hand gesture classification using low-end wearable sensor combined with unsupervised transfer learning, we adopted and implemented the Self-Calibrating Asynchronous Domain Adversarial Neural Network (SCADANN) developed by Côté-Allard et al. [30].

#### 2.2.1. Overview of SCDANN

Detailed description of the framework and algorithms can be found in the original paper [30]. Briefly, SCADANN automatically adapts a deep learning network pre-trained with labeled source data to improve the accuracy for the new, unlabeled target data. This is achieved by three steps involving number of networks and algorithms:

1. Apply Domain Adversarial Neural Network (DANN) [31] to the network using the labeled and newly acquired unlabeled data. Domain adversarial training aims to improve classification performance of the network by learning a feature representation that favors class separability of the labeled dataset, while reducing domain divergence (i.e., differentiation) between the labeled recording session and the unlabeled recording.
2. Using the adapted network, perform relabeling with pseudo-labels generating heuristic. This heuristic builds on Multiple Votes self-recalibration approach proposed by Zhai et al. [32], which leverages the prediction’s context to generate pseudo-labels considering the median softmax value of the network’s output from surrounding segments in recorded data. Côté-Allard et al. enhanced the robustness and efficiency of this recalibration by detecting stable gesture transitions, replacing short-term unstable predictions with pseudo labels, and ignoring long-term unstable readings.
3. Starting from the adapted network, train the network with the pseudo-labeled data and labeled data while continuing to apply DANN to minimize domain divergence. Compared to the first step, here the network’s weights are also optimized in relation to the cross-entropy loss calculated from the newly generated pseudo-labels.

#### 2.2.2. Deep learning-based classification models

To test and compare the performance of SCADANN when applied to different network prediction models (see ***Study design and analyses*** section below), we implemented two deep learning based neural networks (Fig. 2), as in the original work of Côté-Allard et al. [30]:

1. Deep neural network with Temporal-Spatial Descriptors (TSD) [33]: TSD employed a small and simple deep neural network architecture with 3 fully connected layers, each 200 neurons wide and applying batch normalization, leaky ReLU (slope=0.1), and dropout (set to p=0.5). As indicated by its name, handcrafted feature sets for EMG-based gesture classification proposed by Khushaba et al. [33] was used. Specifically, this feature set, i.e., TSD, gives an enhanced representation of muscle activities through incorporating the information that can be extracted from individual EMG channels (signal time-domain descriptors) as well as combined EMG channels (temporal-spatial correlation features). Total 252 features extracted from each segment of the Myo dataset (8 channels) were used as input for TSD; the full list and description of feature set can be found in the original paper [33].
2. Convolutional neural Network with spectrogram (ConvNet) [30]: ConvNet also employed a small and simple convolutional neural network architecture, which contained four blocks (each encapsulating a convolutional layer, batch normalization, leaky ReLU with slope 0.1, and dropout set to p=0.5) followed by a global average polling and two heads (gesture and domain output). This architecture was originally inspired by [34]. Spectrograms of all 8 channels for a given segment, 4 × 8 × 10 (time × channel × frequency) with 53,528 learnable parameters, were used as input for the ConvNet.

**Figure 2.**
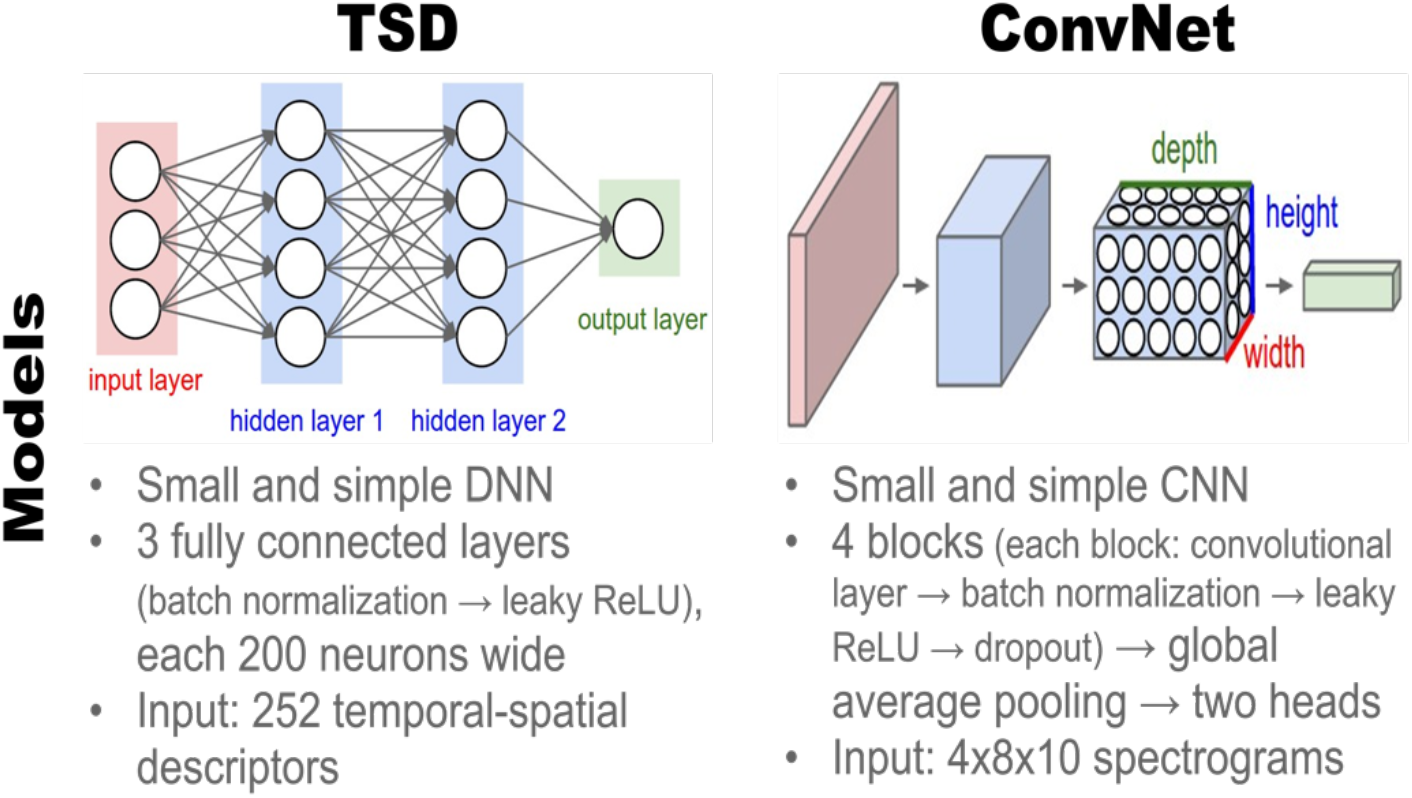
Deep learning models. Schematics of the two deep learning-based gesture classification models employed: Deep neural network with Temporal-Spatial Descriptors (TSD, left) [33] and Convolutional neural Network with spectrogram (ConvNet, right) [30].

#### 2.2.3. Implementation for Myo Dataset

The original dataset used to develop and validate the SCADANN algorithm was acquired by a laboratory built wearable device 3DC Armband [35], which has a similar form factor but with higher technical specification compared to the Myo Armband: 2.4 GHz wireless low-power consumption protocol, 10-channel (×1.25 greater number) dry-electrode, 1000 Hz sampling rate (×5 greater bandwidth in EMG spectra), 10 bit analog-to-digital conversion (×1.25 greater resolution). The data consisted of EMG recordings from young healthy participants repeatedly performing eleven gestures in three separate sessions over a period of fourteen days (i.e., seven days interval). This data was used to evaluate the performance of SCADANN in preserving gesture classification accuracy across measurement days, in which a small electrode shift and inter-day variability was expected to occur.

Because our goal was to test the feasibility of applying unsupervised transfer learning framework to a lower-grade device in more variable contexts (inter-user, -day, and -location) in most conservative manner, we did not optimize any model or algorithm hyperparameters (e.g., learning rate). All algorithms (SCADANN, TSD, and ConvNet) were appropriately modified only to take the Myo dataset as input, which differed in dimensions, bandwidth, and resolution from the original dataset as described above. All algorithms were written in Python, implemented with PyTorch library and are made available, together with data supporting the results reported in this study, in public repository: https://github.com/aonai/long_term_EMG_myo.

### 2.3. Study design and analyses

#### 2.3.1. Inter-user, -day, and -location transfer learning

Our main goal was to systematically evaluate the performance of the proposed transfer learning (SCADANN) in improving prediction accuracy across various measuring conditions in the Myo dataset: 5 users × 10 days × 3 sensor locations. To this end, we first trained a deep neural network (TSD) with labeled source data from any particular user, day, or sensor location, then applied the SCADANN for unlabeled target data from a different user, day, or location. Specifically, we employed a design for selecting source and target data in which the effect of transferring across each of the three conditions can be best evaluated in isolation (Fig. 3). For inter-user transfer, every possible User-User pair was examined, separately for each of the three locations (N, I, O) whereas data across 10 days in the same location were lumped. For inter-day transfer, due to too large number of possible pairs, Day 1 was used as source data and Day 2, 3, …, 10 as target data, separately for each user and each location. For inter-location transfer, Location N was used as source data and other two locations (I and O) as target data, separately for each user whereas data across 10 days in the same location were lumped.

**Figure 3.**
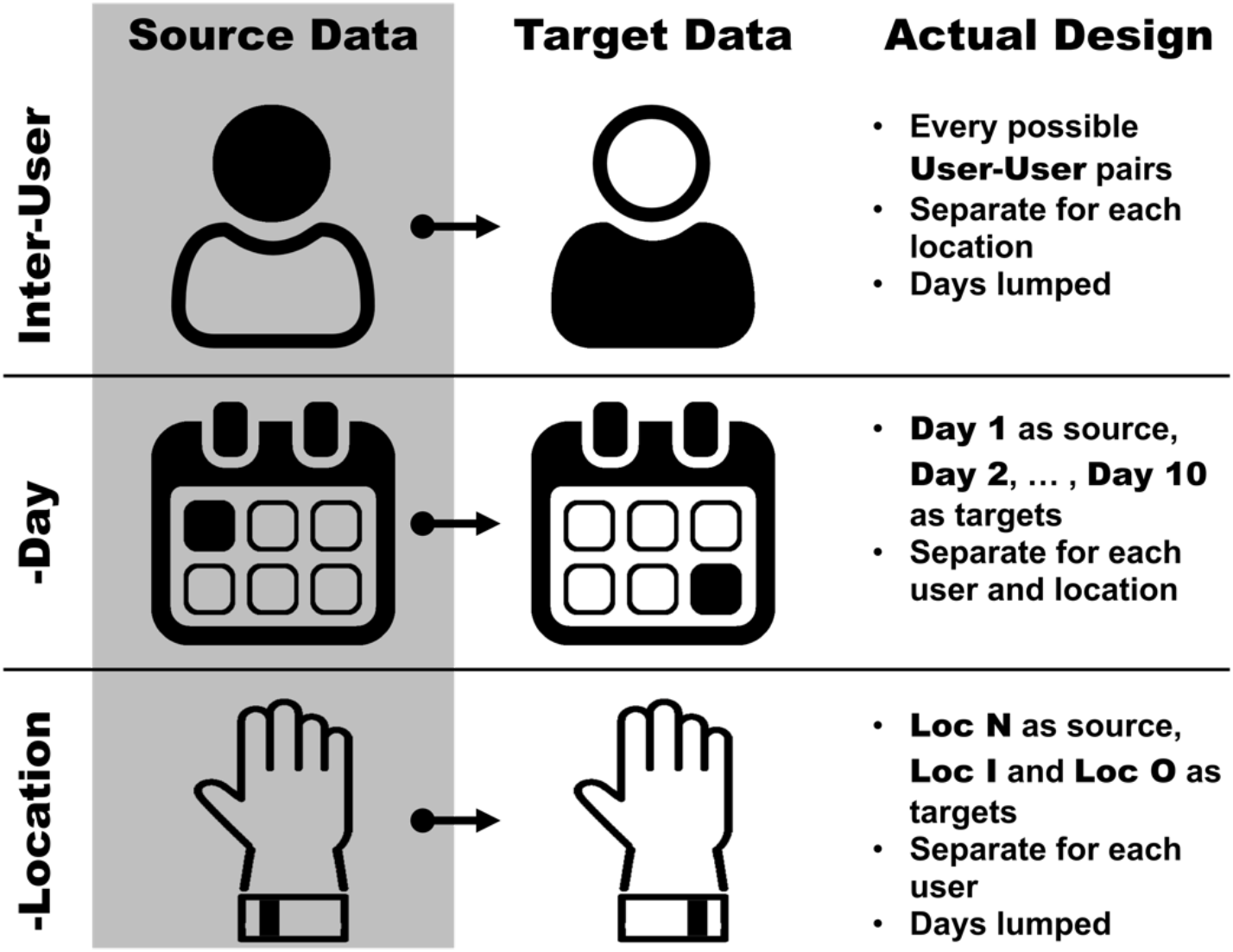
Design for evaluating performance of transfer learning. TSD was trained with labeled source data from any particular user, day, or location. Then, the SCADANN was applied for unlabeled target data from a different user, day, or location.

Performance of transfer learning was evaluated with improvement in classification accuracy after applying SCADANN, compared to when the original TSD was directly applied to target data without recalibration. Classification accuracy was calculated offline, defined as the percentage of time the network predicted the correct gesture compared with ground truth, over total 26 samples in each 1.5 s segment data (250 ms window with 80% overlap) times the number of trials lumped. While accuracy was computed for each gesture in each trial, we report results averaged across four repetitions, across all 22 gestures, and across the total number of data lumped depending on the transferring condition (Fig. 3), which will be referred to as transfer accuracy.

In an attempt to elucidate the informative context in which SCADANN performs better or worse, we further examined the correlation between the cross-over accuracy of the original TSD, i.e., with No Recalibration (TSD NoRecal), and transfer accuracy of the SCADANN. A Pearson correlation coefficient was calculated for averaged accuracy in each transfer case (described above), which resulted in 3, 3, and 2 cases for each of the five users in the inter-user (total 15 cases), inter-day (total 15 cases), and inter-location (10 cases) transfer cases, respectively. Results indicated strong correlations in all cases (see section 3), which led us to further examine additional contexts in which performance of the SCADANN can be improved, as described below.

#### 2.3.2. Effect of initial model selection (TSD vs. ConvNet)

While SCADANN can be applied to automatically recalibrate any prediction model – most ideally, deep learning networks [30], different models can yield different performance initially when being trained with source data. Despite that initially poor performance (accuracy) of the original model may leave more room for improvement, domain adversarial training inherently is affected by the distribution and (potentially hidden) feature representation within the labeled dataset in the source domain. Thus, it is likely that the model with initially poor performance will also yield degraded performance after transfer learning. To investigate the effect of initial model selection, we compared the performance of SCDANN when applied to two different models TSD and ConvNet described above, examined in one example case of inter-location transfer using data from one representative user (User 1), lumped over 10 days. Classification accuracy of both the original model trained with the same source data and after applying SCADANN were compared.

#### 2.3.3. Effect of incremental source data

While the main purpose of transfer learning is to minimize the need for re-calibration or training case-specific models, limited amount of source data or source data with less variability (i.e., not covering potential variation in target data) will negatively impact the performance of transfer learning [8, 27]. One solution to address this issue is to incrementally include more source data, for example, from more users or over time [36, 37], where “ideal” (minimal and/or best) combination of source data may suffice to ensure robust performance without the need to re-calibrate. To investigate the effect of incremental source data, we evaluated the performance of SCADANN applied to TSD, while cumulatively including data from more users and days in the source data for example inter-user (at location N, days lumped) and inter-day (User 5, at location I) cases, respectively.

## 3. RESULTS

### 3.1. Inter-user, -day, and -location transfer learning

Accuracy was improved in all inter-user, -day, and -location transfers with SCADANN applied to a TSD model initially trained with a particular user, day, and sensor location (source data) to predict gestures in a different user, day, or sensor location (target data). For example, for inter-user transfer, TSD initially achieved accuracy of 87.0% and 70.1% for User 1 and User 3, respectively, on the data from each individual lumped across 10 days with a sensor worn at the neutral location (Fig. 4, top left TSD Original in filled square). When the TSD trained with this source data was applied to target data from other users (i.e., each User 2, 3, and 4 for User 1 and User 1, 2, 4 for User 3) without recalibration, accuracy was substantially decreased in all cases (Fig.4, top left; TSD NoRecal in dotted line). After applying SCADANN, accuracy improved in all cases (Fig.4, top left, SCADANN in solid line), with transfer from User 1 to other users generally achieving higher accuracy than transfer from User 3. Similarly, for inter-day and inter-location transfer, the degraded accuracy of the original TSD without recalibration could be substantially improved (Fig.4, top middle and right, respectively). Overall, SCADANN improved accuracy by 12±5.2% (avg±sd), 9.6±5.0%, and 9.3±3.5% across all user-to-user, day-to-day, and location-to-location cases, respectively (Fig.4, middle row).

**Figure 4.**
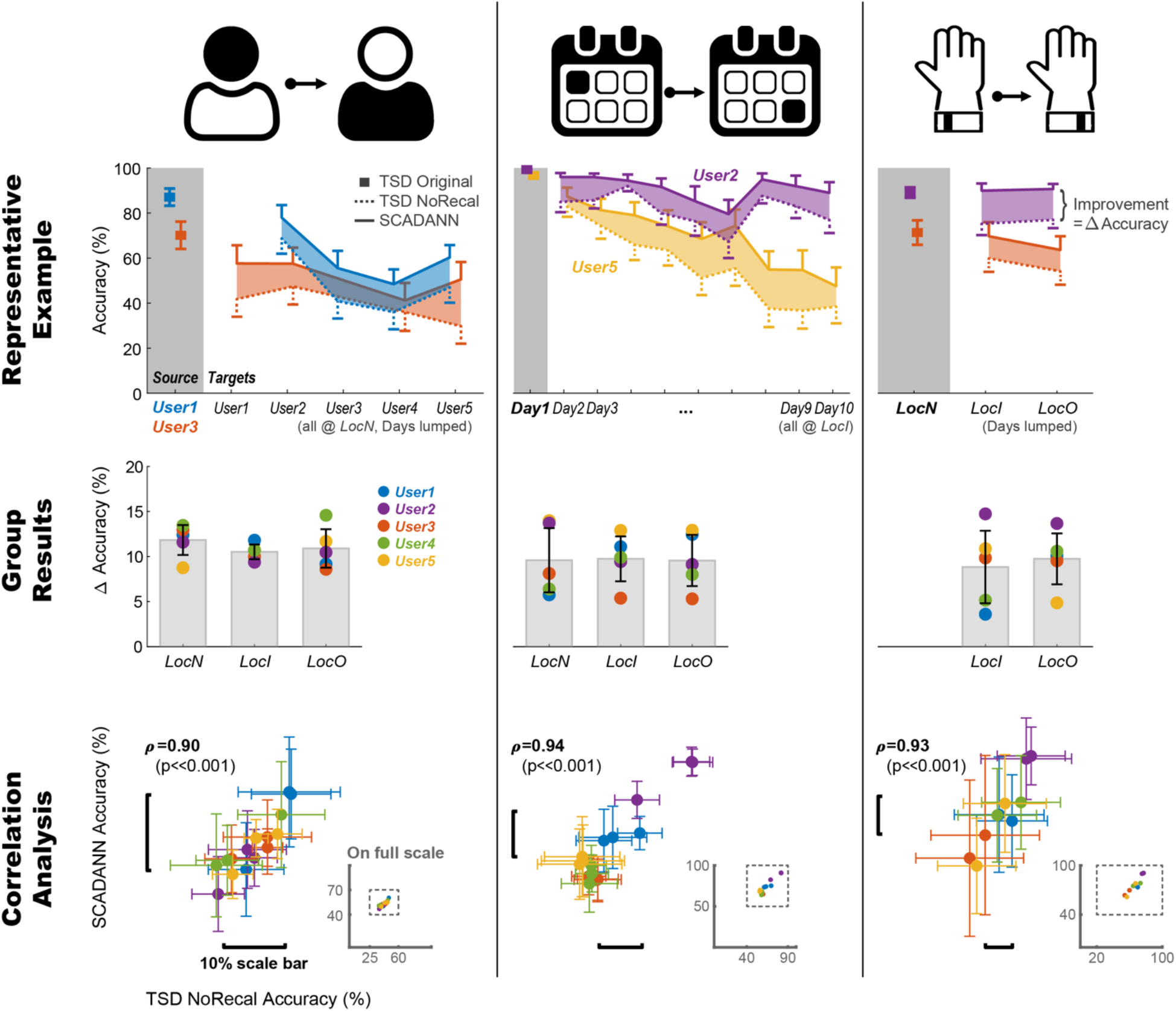
Transfer learning results. Accuracy after applying SCDANN (solid line), compared to the original TSD with no recalibration (dotted line), in the case of inter-user (left column), -day (middle column), and -location (right column) transfer; color coded by user. Top and middle row show representative examples and group results for each transfer case, respectively. Bottom row shows correlation analysis between the cross-over accuracy of the original TSD vs. the transfer accuracy of the SCADANN; insets show the same data on full scale (0-100%).

As seen from representative examples (Fig.4, top row), the amount of improvement varied depending on cases. In one best-case scenario, accuracy improved by 26% (from 67% to 93%), whereas sometimes the gain is modest (e.g., from 76% to 78%). In general, cases with higher cross-over accuracy of the original TSD, (i.e., without recalibration), also achieved greater transfer accuracy after applying SCADANN. For example, User 2. vs. User 5 in inter-day transfer (Fig. 4 top middle) or User 2 vs. User 3 in inter-day transfer (Fig. 4 top right). This observation was confirmed with correlation analysis, where we found strong correlation (rho>0.90, p<<0.001) between the cross-over accuracy of the original TSD without recalibration and transfer accuracy of the SCADANN in all inter-user, -day, and -location transfers (Fig. 4 bottom row).

### 3.2. Effect of initial model selection (TSD vs. ConvNet)

We found that classification accuracy of both the original model (i.e., for source data) and after transfer (i.e., applying to target with SCADANN) were greater for TSD, a simple deep learning model with Temporal-Spatial Descriptors [33], compared to ConvNet, a simple convolutional neural network with spectrogram [30]. For the one inter-location transfer case example examined, using data from User 1, lumped over 10 days, classification accuracy was greater in TSD (95.2% on average) compared to ConvNet (79.2% on average) for the source data, i.e., Location N (Fig. 5, left). The accuracy of TSD was greater than ConvNet for most of the 22 gestures except for one 2-DoF gesture, in which accuracy of ConvNet was greater than TSD by only 2%.

**Figure 5.**
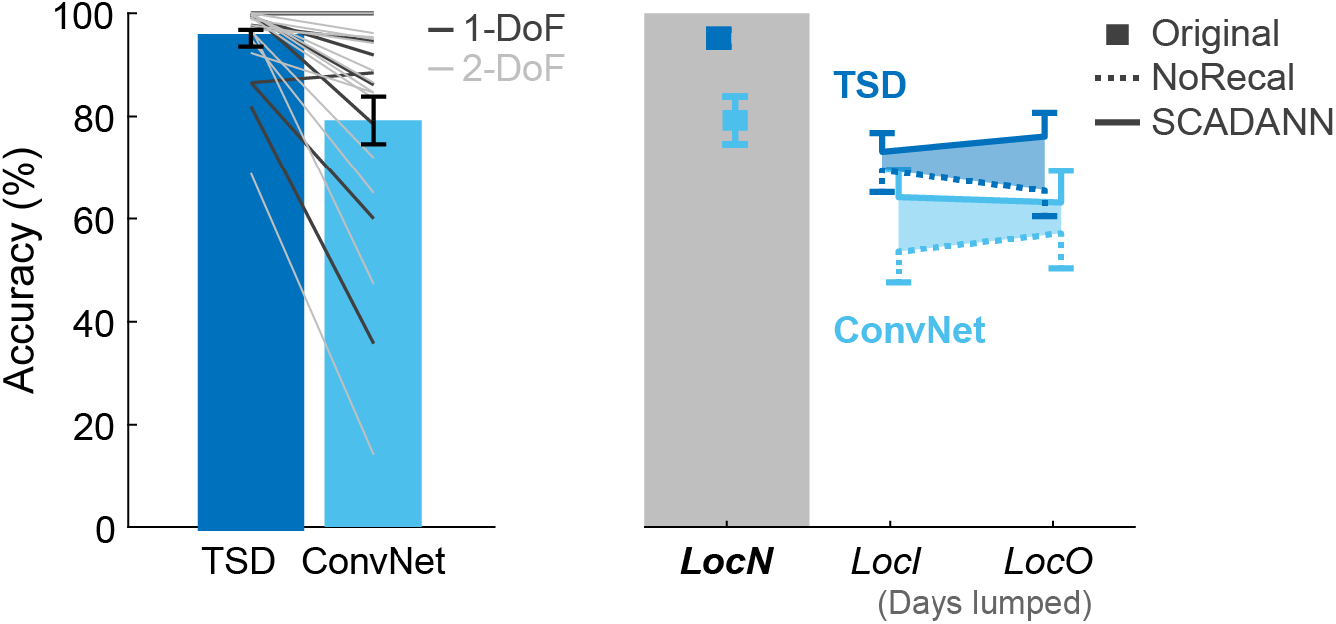
Effect of initial model selection. Representative example for inter-location transfer, comparing cross-over accuracy (left) and transfer accuracy (right) of TSD vs. ConvNet.

When applied to target data, i.e., Location I and Location O, accuracy of both TSD and ConvNet was lower, as expected, than that of the original model. Nevertheless, TSD maintained higher crossover accuracy compared to ConvNet for both locations (Fig. 5, right, NoRecal with dotted line). Furthermore, transfer accuracy after applying SCADANN was higher for TSD compared to ConvNet for both locations.

While we only examined one case as a representative example, the substantially better performance described above and the very strong correlation between the cross-over accuracy and the transfer accuracy (Fig. 4, bottom row) suggest that TSD will outperform ConvNet in most of the cases, including inter-day and inter-user transfers.

### 3.3. Effect of incremental source data

We also found that cumulatively including data from more users or days in the source data tends to improve performance for new target data. For the one inter-user transfer case example examined (Fig. 6, top), using incremental source data from User 1, User 1+2, User 1+2+3, and User 1+2+3+4, lumped over 10 days for sensor worn at Location N, transfer accuracy improved compared to using only User 1 as source data in three out of five cases (5.70% greater accuracy on average). In the two cases, transfer accuracy was slightly lower (1.55% on average).

**Figure 6.**
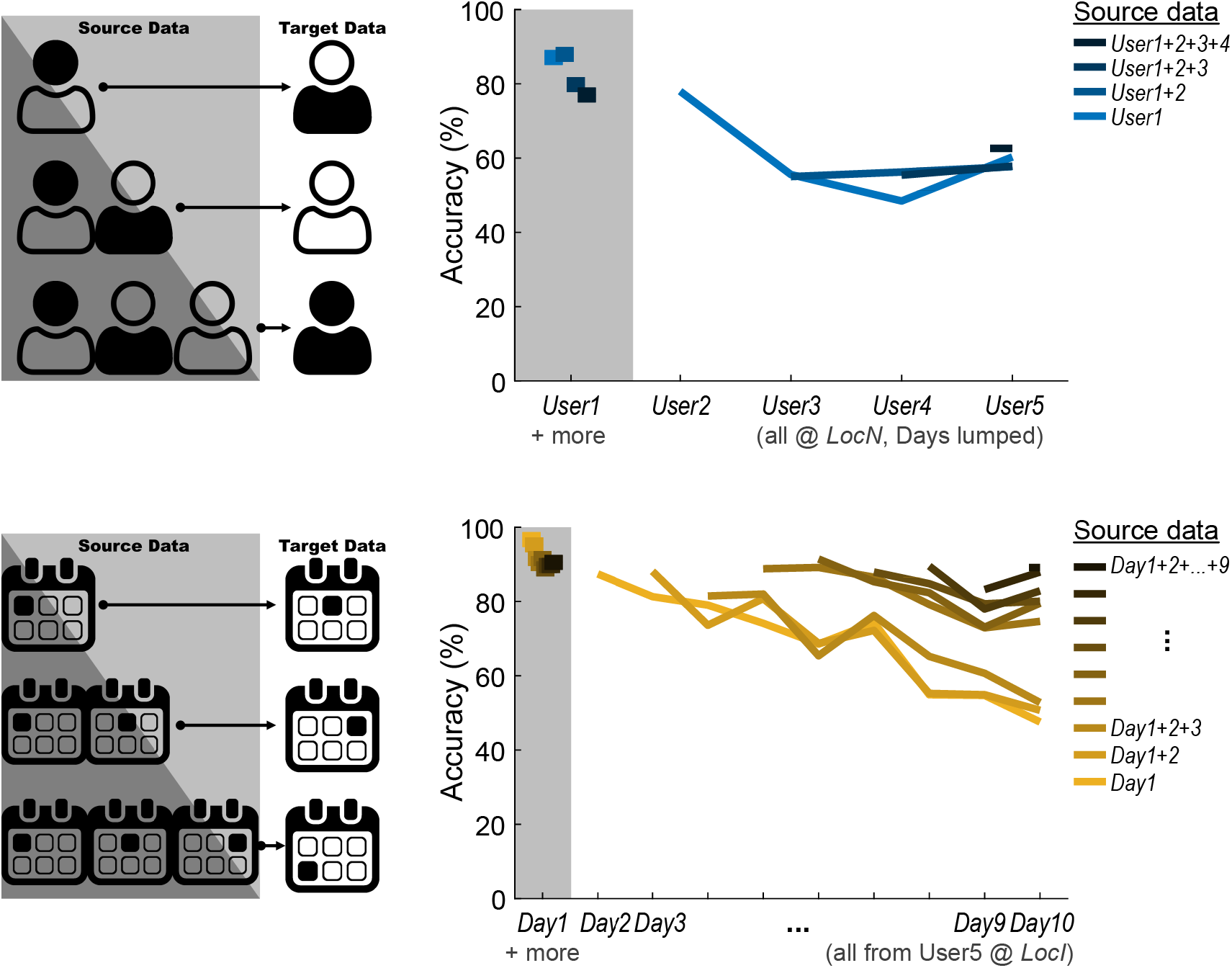
Effect of incremental source data. Representative examples for inter-user (top) and inter-day (bottom) transfer learning.

On the other hand, for the one inter-day transfer case examined (Fig. 6, bottom), using incremental source data from Day 1, Day 1+2, …, Day 1+2+…+9 of User 5 with sensor worn at Location I improved transfer accuracy compared to using only Day 1 as source data in most cases (17.7% greater on average in 33 out of 36 cases; 3.55% lower on average in 3 cases). Especially, transfer accuracy was substantially improved (25.4% on average, 41.6% maximum) after including Day 1 to Day 4 as source data compared to single day (Day 1) and maintained relatively high accuracy. While transfer accuracy continued to improve as data from more days were included, data beyond Day 4 however resulted in a modest gain, indicating that SCADANN is most efficient when recording from the first four days are selected as the source data.

## 4. DISCUSSION

Our goal was to test the feasibility of using a low-end wearable EMG sensor and unsupervised transfer learning for robust and seamless myoelectric control. Using a hand gesture classification dataset acquired with a consumer-grade sensor (Myo Armband) and a domain adversarial adaptation algorithm SCADANN [30], we demonstrate that transfer learning can improve accuracy across various measurement conditions, without the need to recalibrate. Specifically, a deep learning model (TSD) trained with source data (i.e., particular user, day, or location) could predict unlabeled target data from a different user, day, or sensor location with improved accuracy. We also found the performance of the original model (TSD) without recalibration dictates performance of transfer learning (SCADANN), which implies that a “good” model trained with “good” source data would yield better transfer accuracy. Indeed, representative examples illustrated that better selection of initial model (e.g., TSD vs. ConvNet) and training that model with incremental source data could further improve performance of transfer learning.

This work is the first, to our knowledge, to systematically investigate the feasibility of using low-end wearable EMG sensor with unsupervised transfer learning while isolating the effect of various factors that contribute to poor cross-over performance without recalibration. Transfer learning applied to myoelectric control has largely focused on inter-user, -day, or -location transfers separately [26–28], and not necessarily explicitly controlling for other factors. Our design, in contrast, allowed us to evaluate each factor independently while controlling for others, by proper selection and combination of source and targe data available from the dataset. Some studies that controlled for more than two factors have used supervised or semi-supervised approaches in which the at least some of the target data was labeled [37–39]. On the other hand, other unsupervised transfer learning frameworks have been developed and validated with data acquired using sensors with better technical specifications such as a faster sampling rate and more channels [20, 35], both of which tends to improve performance [20]. The virtue of using a low-end sensor combined with unsupervised transfer learning is that it increases acceptability and usability, without needing to retrain the model, or, for example, needing to acquire new labeled target data.

Another contribution of this work is that we further provide informative contexts in which the performance of transfer learning can be improved. Both from correlation analysis across all transfer cases extracted from the dataset and specific model comparison (TSD vs. ConvNet), we showed that the model with better initial cross-over accuracy (i.e., with no recalibration) yields better transfer accuracy, which is in line with other studies [30, 37]. We also showed that incremental source data, that is, cumulatively including more data as source for training, tends to improve performance. Interestingly, however, more data was not always beneficial, suggesting that to obtain the best outcome, one should find a balance between an “ideal” combination of only a few datasets and the resulting performance. Indeed, source data selection is an important problem in transfer learning [8, 27], which warrants future investigation. For example, the analysis approach taken in this work provides a framework in which one can systematically assess source data selection, as well as combine the effects of transfer learning across contexts (e.g., user × day) using layers of domain adversarial networks [40].

The main limitation of this work is that because our goal was to demonstrate the feasibility of using low-end wearable EMG sensor and unsupervised transfer learning, we did not explicitly and rigorously fine-tune or validate the employed dataset and algorithms. Nevertheless, we postulate the proposed approach is feasible and promising, given that our results achieved comparable performance (i.e., improvement in accuracy) to similar approaches to date [30, 37], even without data- or algorithm-specific parameter optimization and with a limited dataset (e.g., only five users). The proposed approach, further tailored for specific tasks or applications will confer more robust performance of unsupervised transfer learning. Furthermore, the framework employed in this study can be readily extended to more advanced myoelectric control applications, such as for more complex or dynamic human behavior, e.g., especially when integrated with combined EMG and inertial measurement unit data [41], as well as online or real-time control requiring minimal delay [42–45].

## 5. CONCLUSIONS

We propose that robust yet seamless myoelectric control can be achieved using a low-end, easy-to-“don” and “doff” wearable EMG electrodes combined with unsupervised transfer learning. In this study, we demonstrate that such approach is feasible and promising by systematically investigating whether, and in what contexts unsupervised learning can be applied to improve performance of gesture classification using data from a wearable EMG armband across various measurement conditions such as users, sensor locations, and times. The proposed framework can be readily applied for human-machine interface applications with myoelectric control such as assistive and/or rehabilitative robotic devices and virtual reality.

## Author Contributions

Conceptualization, M.H.S.; methodology, M.H.S. and S.Y.L.; software, S.Y.L.; validation, M.H.S. and S.Y.L.; formal analysis, M.H.S. and S.Y.L.; investigation, M.H.S. and S.Y.L.; resources, M.L.E. and J.P.A.D.; data curation, M.H.S. and S.Y.L.; writing—original draft preparation, M.H.S. and S.Y.L.; writing—review and editing, M.H.S., S.Y.L., M.L.E. and J.P.A.D.; visualization, M.H.S.; supervision, M.H.S., M.L.E. and J.P.A.D.; project administration, M.L.E.; funding acquisition, M.L.E. and J.P.A.D. All authors have read and agreed to the published version of the manuscript.

## Institutional Review Board Statement

Not applicable.

## Data Availability Statement

All data and codes used in this paper are available on public repository: https://github.com/aonai/long_term_EMG_myo.

## Acknowledgments

Any support given which is not covered by the author contribution or funding sections, e.g., administrative and technical support, or donations in kind.

## REFERENCES

1. Oskoei, M. A., and H. S. Hu. “Myoelectric Control Systems-a Survey.” Biomedical Signal Processing and Control 2, no. 4 (2007): 275–94.

2. Fraser, G. D., A. D. C. Chan, J. R. Green, and D. T. MacIsaac. “Automated Biosignal Quality Analysis for Electromyography Using a One-Class Support Vector Machine.” Ieee Transactions on Instrumentation and Measurement 63, no. 12 (2014): 2919–30.

3. Muceli, S., and D. Farina. “Simultaneous and Proportional Estimation of Hand Kinematics from Emg During Mirrored Movements at Multiple Degrees-of-Freedom.” Ieee Transactions on Neural Systems and Rehabilitation Engineering 20, no. 3 (2012): 371–78.

4. Scheme, E., and K. Englehart. “Electromyogram Pattern Recognition for Control of Powered Upper-Limb Prostheses: State of the Art and Challenges for Clinical Use.” Journal of Rehabilitation Research and Development 48, no. 6 (2011): 643–59.

5. Guo, Y., S. Gok, and M. Sahin. “Convolutional Networks Outperform Linear Decoders in Predicting Emg from Spinal Cord Signals.” Front Neurosci 12 (2018): 689.

6. Ketyko, I., F. Kovacs, and K. Z. Varga. “Domain Adaptation for Semg-Based Gesture Recognition with Recurrent Neural Networks.” 2019 International Joint Conference on Neural Networks (Ijcnn) (2019).

7. Ferris, D. P., and C. L. Lewis. “Robotic Lower Limb Exoskeletons Using Proportional Myoelectric Control.” Annu Int Conf IEEE Eng Med Biol Soc 2009 (2009): 2119–24.

8. Yao, Y., and G. Doretto. “Boosting for Transfer Learning with Multiple Sources.” 2010 Ieee Conference on Computer Vision and Pattern Recognition (Cvpr) (2010): 1855–62.

9. Allard, U. C., F. Nougarou, C. L. Fall, P. Giguere, C. Gosselin, F. Laviolette, and B. Gosselin. “A Convolutional Neural Network for Robotic Arm Guidance Using Semg Based Frequency-Features.” 2016 Ieee/Rsj International Conference on Intelligent Robots and Systems (Iros 2016) (2016): 2464–70.

10. Hassan, Hussein F., Sadiq J. Abou-Loukh, and Ibraheem Kasim Ibraheem. “Teleoperated Robotic Arm Movement Using Electromyography Signal with Wearable Myo Armband.” Journal of King Saud University - Engineering Sciences 32, no. 6 (2020): 378–87.

11. Meattini, R., S. Benatti, U. Scarcia, D. De Gregorio, L. Benini, and C. Melchiorri. “An Semg-Based Human-Robot Interface for Robotic Hands Using Machine Learning and Synergies.” Ieee Transactions on Components Packaging and Manufacturing Technology 8, no. 7 (2018): 1149–58.

12. Cote-Allard, U., D. St-Onge, P. Giguere, F. Laviolette, and B. Gosselin. “Towards the Use of Consumer-Grade Electromyographic Armbands for Interactive, Artistic Robotics Performances.” 2017 26th Ieee International Symposium on Robot and Human Interactive Communication (Ro-Man) (2017): 1030–36.

13. Li, Xiaoou, Zhiyong Zhou, Wanyang Liu, and Mengjie Ji. “Wireless Semg-Based Identification in a Virtual Reality Environment.” Microelectronics Reliability 98 (2019): 78–85.

14. Woodward, R. B., and L. J. Hargrove. “Adapting Myoelectric Control in Real-Time Using a Virtual Environment.” J Neuroeng Rehabil 16, no. 1 (2019): 11.

15. Roy, S. H., G. De Luca, M. S. Cheng, A. Johansson, L. D. Gilmore, and C. J. De Luca. “Electro-Mechanical Stability of Surface Emg Sensors.” Medical & Biological Engineering & Computing 45, no. 5 (2007): 447–57.

16. Beck, T. W., T. J. Housh, J. T. Cramer, M. H. Malek, M. Mielke, R. Hendrix, and J. P. Weir. “Sensor Shift and Normalization Reduce the Innervation Zone’s Influence on Emg.” Medicine and Science in Sports and Exercise 40, no. 7 (2008): 1314–22.

17. Hakonen, M., H. Piitulainen, and A. Visala. “Current State of Digital Signal Processing in Myoelectric Interfaces and Related Applications.” Biomedical Signal Processing and Control 18 (2015): 334–59.

18. He, J. Y., D. G. Zhang, N. Jiang, X. J. Sheng, D. Farina, and X. Y. Zhu. “User Adaptation in Long-Term, Open-Loop Myoelectric Training: Implications for Emg Pattern Recognition in Prosthesis Control.” Journal of Neural Engineering 12, no. 4 (2015).

19. Stegeman, D. F., B. U. Kleine, B. G. Lapatki, and J. P. Van Dijk. “High-Density Surface Emg: Techniques and Applications at a Motor Unit Level.” Biocybernetics and Biomedical Engineering 32, no. 3 (2012): 3–27.

20. Pizzolato, S., L. Tagliapietra, M. Cognolato, M. Reggiani, H. Muller, and M. Atzori. “Comparison of Six Electromyography Acquisition Setups on Hand Movement Classification Tasks.” Plos One 12, no. 10 (2017).

21. Feldner, H. A., D. Howell, V. E. Kelly, S. W. McCoy, and K. M. Steele. ““Look, Your Muscles Are Firing!”: A Qualitative Study of Clinician Perspectives on the Use of Surface Electromyography in Neurorehabilitation.” Archives of Physical Medicine and Rehabilitation 100, no. 4 (2019): 663–75.

22. Feldner, H. A., C. Papazian, K. Peters, and K. M. Steele. ““It’s All Sort of Cool and Interesting…But What Do I Do with It?” A Qualitative Study of Stroke Survivors’ Perceptions of Surface Electromyography.” Front Neurol 11 (2020): 1037.

23. Tran, V. T., C. Riveros, and P. Ravaud. “Patients’ Views of Wearable Devices and Ai in Healthcare: Findings from the Compare E-Cohort.” NPJ Digit Med 2 (2019): 53.

24. Prahm, Cosima, Benjamin Paassen, Alexander Schulz, Barbara Hammer, and Oskar Aszmann. “Transfer Learning for Rapid Re-Calibration of a Myoelectric Prosthesis after Sensor Shift.” In Converging Clinical and Engineering Research on Neurorehabilitation Ii, 153–57, 2017.

25. Motiian, S., M. Piccirilli, D. A. Adjeroh, and G. Doretto. “Unified Deep Supervised Domain Adaptation and Generalization.” 2017 Ieee International Conference on Computer Vision (Iccv) (2017): 5716–26.

26. Prahm, C., A. Schulz, B. Paaben, J. Schoisswohl, E. Kaniusas, G. Dorffner, B. Hammer, and O. Aszmann. “Counteracting Sensor Shifts in Upper-Limb Prosthesis Control Via Transfer Learning.” IEEE Trans Neural Syst Rehabil Eng 27, no. 5 (2019): 956–62.

27. Wei, C. S., Y. P. Lin, Y. T. Wang, C. T. Lin, and T. P. Jung. “A Subject-Transfer Framework for Obviating Inter- and Intra-Subject Variability in Eeg-Based Drowsiness Detection.” Neuroimage 174 (2018): 407–19.

28. Zia Ur Rehman, M., A. Waris, S. O. Gilani, M. Jochumsen, I. K. Niazi, M. Jamil, D. Farina, and E. N. Kamavuako. “Multiday Emg-Based Classification of Hand Motions with Deep Learning Techniques.” Sensors (Basel) 18, no. 8 (2018).

29. Kanoga, Suguru, Atsunori Kanemura, and Hideki Asoh. “Are Armband Semg Devices Dense Enough for Long-Term Use?—Sensor Placement Shifts Cause Significant Reduction in Recognition Accuracy.” Biomedical Signal Processing and Control 60 (2020).

30. Cote-Allard, Ulysse, Gabriel Gagnon-Turcotte, Angkoon Phinyomark, Kyrre Glette, Erik J. Scheme, Francois Laviolette, and Benoit Gosselin. “Unsupervised Domain Adversarial Self-Calibration for Electromyography-Based Gesture Recognition.” IEEE Access 8 (2020): 177941–55.

31. Ganin, Y., E. Ustinova, H. Ajakan, P. Germain, H. Larochelle, F. Laviolette, M. Marchand, and V. Lempitsky. “Domain-Adversarial Training of Neural Networks.” Journal of Machine Learning Research 17 (2016).

32. Zhai, X. L., B. Jelfs, R. H. M. Chan, and C. Tin. “Self-Recalibrating Surface Emg Pattern Recognition for Neuroprosthesis Control Based on Convolutional Neural Network.” Frontiers in Neuroscience 11 (2017).

33. Khushaba, R. N., A. H. Al-Timemy, A. Al-Ani, and A. Al-Jumaily. “A Framework of Temporal-Spatial Descriptors-Based Feature Extraction for Improved Myoelectric Pattern Recognition.” Ieee Transactions on Neural Systems and Rehabilitation Engineering 25, no. 10 (2017): 1821–31.

34. Cote-Allard, U., E. Campbell, A. Phinyomark, F. Laviolette, B. Gosselin, and E. Scheme. “Interpreting Deep Learning Features for Myoelectric Control: A Comparison with Handcrafted Features.” Frontiers in Bioengineering and Biotechnology 8 (2020).

35. Cote-Allard, U., G. Gagnon-Turcotte, F. Laviolette, and B. Gosselin. “A Low-Cost, Wireless, 3-D-Printed Custom Armband for Semg Hand Gesture Recognition.” Sensors 19, no. 12 (2019).

36. Gallego, A. J., J. Calvo-Zaragoza, and R. B. Fisher. “Incremental Unsupervised Domain-Adversarial Training of Neural Networks.” Ieee Transactions on Neural Networks and Learning Systems 32, no. 11 (2021): 4864–78.

37. Kanoga, S., T. Hoshino, and H. Asoh. “Semi-Supervised Style Transfer Mapping-Based Framework for Semg-Based Pattern Recognition with 1-or 2-Dof Forearm Motions.” Biomedical Signal Processing and Control 68 (2021).

38. Schwarz, A., J. Brandstetter, J. Pereira, and G. R. Muller-Putz. “Direct Comparison of Supervised and Semi-Supervised Retraining Approaches for Co-Adaptive Bcis.” Medical & Biological Engineering & Computing 57, no. 11 (2019): 2347–57.

39. Vidovic, M. M. C., H. J. Hwang, S. Amsuss, J. M. Hahne, D. Farina, and K. R. Muller. “Improving the Robustness of Myoelectric Pattern Recognition for Upper Limb Prostheses by Covariate Shift Adaptation.” Ieee Transactions on Neural Systems and Rehabilitation Engineering 24, no. 9 (2016): 961–70.

40. Gu, X., Y. Guo, F. Deligianni, B. Lo, and G. Z. Yang. “Cross-Subject and Cross-Modal Transfer for Generalized Abnormal Gait Pattern Recognition.” IEEE Trans Neural Netw Learn Syst PP (2020).

41. Maceira-Elvira, P., T. Popa, A. C. Schmid, and F. C. Hummel. “Wearable Technology in Stroke Rehabilitation: Towards Improved Diagnosis and Treatment of Upper-Limb Motor Impairment.” J Neuroeng Rehabil 16, no. 1 (2019): 142.

42. Hu, Xuhui, Hong Zeng, Dapeng Chen, Jiahang Zhu, and Aiguo Song. “Real-Time Continuous Hand Motion Myoelectric Decoding by Automated Data Labeling.” (2019).

43. Jaramillo-Yanez, A., M. E. Benalcazar, and E. Mena-Maldonado. “Real-Time Hand Gesture Recognition Using Surface Electromyography and Machine Learning: A Systematic Literature Review.” Sensors (Basel) 20, no. 9 (2020).

44. Parajuli, N., N. Sreenivasan, P. Bifulco, M. Cesarelli, S. Savino, V. Niola, D. Esposito, T. J. Hamilton, G. R. Naik, U. Gunawardana, and G. D. Gargiulo. “Real-Time Emg Based Pattern Recognition Control for Hand Prostheses: A Review on Existing Methods, Challenges and Future Implementation.” Sensors (Basel) 19, no. 20 (2019).

45. Zhang, Z., K. Yang, J. Qian, and L. Zhang. “Real-Time Surface Emg Pattern Recognition for Hand Gestures Based on an Artificial Neural Network.” Sensors (Basel) 19, no. 14 (2019).

